# Body temperature rebounds on sea ice and is elevated by mercury contamination in a keystone predator

**DOI:** 10.1101/2022.10.18.512690

**Authors:** Melissa L. Grunst, Andrea S. Grunst, David Grémillet, Akiko Sato, Sophie Gentès, Jérôme Fort

**Affiliations:** Littoral, Environnement et Sociétés (LIENSs), UMR 7266 CNRS-La Rochelle Université, 2 Rue Olympe de Gouges, FR-17000 La Rochelle, France; CEFE, UMR 5175, CNRS – Université de Montpellier – Université Paul-Valéry Montpellier – EPHE; Montpellier, France; Percy FitzPatrick Institute of African Ornithology, University of Cape Town, Rondebosch, South Africa; Centre d’Etudes Biologiques de Chizé (CEBC), UMR 7372 CNRS-La Rochelle Université, 79360 Villiers-en-Bois, France

**Keywords:** Body temperature, behavior, plasticity, environmental variation, climate change, ecotoxicology, mercury, biologging

## Abstract

Despite overall stability, plasticity in endothermic body temperature (T_b_) occurs, which may facilitate maintenance of crucial activities in the face of climate change-related environmental variations. However, this plasticity may be limited by physiological or energetic constraints, which are potentially exacerbated by other environmental stressors. For instance, chemical contamination may elevate energetic costs and have endocrine disrupting effects that undermine thermoregulation. We leveraged advanced biologging techniques to elucidate how T_b_ varies with different behavioral states and environmental conditions in a keystone Arctic seabird, the little auk (*Alle alle*). We additionally evaluated whether mercury (Hg) contamination independently affected T_b_, or limited or increased state-dependent changes in T_b_. T_b_ was highest and relatively invariable when birds were at the colony, and rebounded when birds were resting on sea ice, following declines while foraging (diving) in polar waters. These results suggest that the colony and sea ice function as thermal refuges for little auks. In addition, T_b_ increased with ambient temperature and relative humidity across behavioral states, and increased with wind speed when birds were flying. Little auks with higher Hg levels had higher, less variable, T_b_ across behaviors and environmental contexts, perhaps reflecting increased metabolic rates linked to detoxification costs. Results provide evidence for environment- and contaminant-related effects on T_b_, but not interactive effects between the two, and suggest that loss of sea ice and increased environmental contamination under global change may have serious implications for T_b_ regulation and energy balance.

## Introduction

Endothermic animals are characterized by the ability to tightly regulate body temperature (T_b_) via endogenous heat production (McNab 2002). This capacity is potentiated by a high metabolic rate, and allows endotherms to occupy a breadth of thermal niches, achieve a degree of thermal independence from environmental temperatures (Khaliq et al. 2014), and sustain impressive levels of activity driven by high muscular power output (Crompton et al. 1978; Hedrick and Hillman 2016). However, tightly regulating T_b_ via thermoregulation also has energetic costs. Consequently, many endothermic species allow adaptive fluctuations T_b_ to minimize thermoregulatory costs and maximize energy balance (Angilletta et al. 2010; McKechnie and Wolf 2019). Fluctuations in T_b_ may reflect weather conditions that modify the thermal gradient between the body and the environment and the costs of thermoregulation. For instance, small, overwintering songbirds often allow reductions in T_b_ during periods of inactivity, which lowers the temperature differential between the body and environment, hence reducing heat loss and conserving energy (McKechnie and Lovegrove 2002; Douglas et al. 2017; Stager et al. 2020). Similarly, animals facing very hot ambient environmental conditions can allow T_b_ to rise, thus reducing thermoregulatory costs and conserving water (Gerson et al. 2019a; McKechnie and Wolf 2019; Cooper et al. 2020). In the context of global change, organisms may increasingly face challenging thermal environments, and adaptive phenotypic plasticity in T_b_ could be an important energy-saving mechanism.

In addition, endothermic T_b_ fluctuates not only with external environmental conditions, but also with activity. Although heat dissipation mechanisms act to maintain T_b_ within a safe range, T_b_ often increases as a result of heat production during intense activities, such as flying (Aulie 1971; Torre-Bueno 19776; Tapper et al. 2020). However, when temperatures drop below the thermal neutral zone, heat generated by energy-intensive activities may offset energetic costs of thermoregulation, in which case minimal changes in T_b_ with activity may be observed (Bruinzeel and Piersma 2008; Humphries and Careau 2011; Careau and Garland 2012). Animals adapted to harness activity to neutralize thermoregulatory costs may suffer under climate change scenarios of warming temperatures because heat generated by activity no longer balances thermoregulatory costs, but rather, potentially leads to thermal stress. Consequently, animals may be forced to reduce activities that fulfill essential ecological and social functions, with non-trivial effects on fitness (Tapper et al. 2020). On the other hand, as an energy conservation mechanism, animals may allow T_b_ to fall during long periods of inactivity, especially when ambient temperatures are low (Brodin et al. 2017). Thus, elucidating how climate change-linked increases in environmental temperature will affect endothermic energy balance depends on understanding how T_b_ is regulated according to both environmental conditions and activity patterns. However, it is challenging to simultaneously measure variation in T_b_, activity, and environmental parameters through time in the wild. As a result, comprehensive studies of dynamic T_b_ regulation in free-living animals remain scarce, limiting our ability to predict responses to climate change.

Furthermore, organisms do not face the energetic challenges of climate change in isolation, but in combination with other anthropogenic disturbance factors, such as exposure to chemical contaminants (Jennsen 2006). Chemical contaminants, such as mercury (Hg), have the potential to disrupt adaptive T_b_ regulation in endotherms via a number of mechanisms. For instance, Hg may undermine effective thermoregulation via endocrine disrupting effects (Rice et al. 2014). Notably, Hg has been shown to interfere with the production of thyroid hormones, which play a central role in thermoregulation (Wada et al. 2009). In addition, Hg could affect the adaptive thermal set point by elevating detoxification costs and resting metabolic rate (Calow 1991), which could be associated with higher T_b_. Although little data specific to Hg is available, both hypo- and hyperthermic responses have been observed in response to chemical contamination, with hypothermic responses proposed to reflect an adaptive response linked to declines in chemical toxicity at lower T_b_ (Leon 2008; Noyes et al. 2009).

In this study, we used a suite of advanced techniques to gain insight into the potential effects of climate change and chemical contamination (i.e. Hg) levels on T_b_ regulation and energetic costs in a keystone Arctic seabird, the little auk (or dovekie, *Alle alle*). In the Arctic, Hg is an especially prevalent contaminant that reaches remote polar regions via a repeated process of condensation and evaporation and bioaccumulates in marine food chains (Morel et al. 1998; Albert et al. 2019; AMAP 2021). Dynamics of Hg exposure in Arctic animals is being influenced by climate changes. For instance, increases in Hg exposure may result via release from permafrost and expansion of the low oxygen subsurface zone, in which inorganic Hg in the ocean is converted to highly toxic methylmercury (Jonsson et al. 2022). We combined the use of internal T_b_ loggers, which recorded abdominal temperature as a proxy of T_b_, miniaturized accelerometers that record dynamic body acceleration, allowing classification of activity patterns, and weather station data to gain insight into patterns of weather- and activity-specific T_b_ regulation. In addition, we obtained blood samples to assess total-Hg concentrations in the red blood cells as a means of assessing whether Hg contamination could affect that ability to maintain stable T_b_.

We generated a suite of general and specific predictions based on our knowledge of the behavior, morphology and energetics of our study species. In general, we predicted that environmental conditions and activity would interact to affect mean levels and variation in core T_b_. More specifically, little auks have a morphology that compromises adaptation for diving and flying. This morphology is characterized by a high wing loading, which results in extremely high energetic costs of flight (Fort et al. 2009; Ste-Marie et al. 2022). Thus, we predicted that T_b_ would increase when birds were flying relative to when birds were at rest at the colony, on sea ice, or water surface, and that this increase would be magnified under conditions that reduce heat exchange between the body and environment (i.e. higher temperature, lower wind speed and relative humidity), which could ultimately limit activity under climate change scenarios. In contrast, endothermic animals diving into cold polar waters face a significant thermal challenge due to the high thermal conductance of water (Grémillet et al. 2015; Favilla and Costa 2020). As a result, diving animals often allow T_b_ to fall below normothermic levels, which may facilitate aerobic dive capacity and limit energetic costs of heat loss to the environment (Niizuma et al. 2007; Favilla and Costa 2020). Thus, we predicted that T_b_ would decline over the course of foraging episodes, and would subsequently increase when birds were resting on sea ice. We also recognized the potential that regional heterothermy, that is, variation in peripheral temperatures, especially in the appendages, might buffer changes in core T_b_ during diving, resulting in relative stability (Niizuma et al. 2007; Ponganis et al. 2003). In general, we also predicted that variation in T_b_ might increase in the context of thermal challenge, which in the Arctic is most commonly experienced in the context of cold stress (this might be especially relevant during resting periods at the colony, on sea ice, or on the water), but which could also involve heat stress, especially during energetically-demanding activities (i.e. flight). Finally, we predicted that higher blood Hg levels might affect the adaptive thermal set point and thermoregulatory capacity. Thus, higher Hg levels could be linked to either higher mean T_b_, perhaps reflecting increased metabolic rates to support detoxification costs, or lower T_b_, perhaps reflecting suppression of thyroid hormones (Chastel et al. 2022). In addition, elevated blood Hg could be linked to greater variation in T_b_, especially in the context of thermal stress.

## Methodology

### Study system

We studied a breeding population of little auks situated at Ukaleqarteq (Kap Höegh), East Greenland (70°44′N, 21°35′W). This population has been the subject of intensive research since 2005. Little auks nest in rock crevasses and can be captured and recaptured at or near the nest site, facilitating fitting and retrieval of accelerometers and deployment of T_b_ loggers. Hg levels in little auks at Ukaleqarteq now exceed toxicological thresholds and evidence suggests negative effects of Hg on reproduction (Fort et al. 2014; Carravieri et al. *unpublished*), energetics (Grunst et al. *unpublished*) and adult body condition (Amélineau et al. 2018).

### Deployment of T_b_ loggers and accelerometers

During July 2020, 8 individuals were simultaneous fitted with T_b_ loggers (BodyCap Anipill Core Body Temperature Ingestible Tablet; BMedical; ±1°C), a telemetric system for gastrointestinal temperature recording, and miniaturized accelerometers (Axy 4, Technosmart, 3g), to record dynamic body acceleration and surface temperature. Upon capture, focal birds ingested T_b_ loggers which recorded abdominal temperature (a proxy for T_b_) every minute for periods of 24h. We then remotely downloaded the data from T_b_ loggers via telemetry when the bird was within ∼1 m of the device. Accelerometers were attached to the breast of the bird at the level of the sternum and positioned centrally using Tesa^®^ tape adhered to the feathers. We marked birds with color rings to facilitate identification and recapture within ∼4 days, upon which we retrieved the accelerometer. Deployment dates all fell within 9 days during the mid-late chick rearing phase [range July 22-30]. A weather station erected at the study site documents variation in ambient weather conditions at a frequency of every 1 minute, including temperature, relative humidity and wind speed.

### Analysis of accelerometry and T_b_ data

Accelerometers recorded data at a frequency of 50 Hz (50 readings per second). We used Igor Pro 8.04 (64-bit; WaveMetrics) to classify data on triaxial acceleration into different behavioral states (see details in Grunst et al. *In Review*). In brief, to identify the time birds spent engaged in different behavioral states through time, we used k-clustering analysis applied to acceleration axes, followed by application of a custom-written script, which utilized both the output from the clustering analysis and surface temperature data. The behavior identified were: flying, diving, on the water surface, on ice, and at the colony. We proceeded to determine whether time spent on the water surface was part of an active foraging bout (i.e. an inter-dive interval), or represented time spent resting on the water surface. To this end, we determined the dive bout ending criterion, using R package diveMove (Luque 2007), which applies the methods of Sibley et al. (1990) and Mori et al. (2001) for the identification of behavioral bouts. We used the standard method of classification, based on the absolute duration of the behavioral bouts (i.e. the inter-dive intervals), rather than the sequential difference method. The bout ending criteria derived was 307.1 seconds. Consequentially, we ended diving bouts if the length of time spent on the water surface exceeded this value, and classified these intervals as time spent resting on the water. Time spent resting on the water additionally encompassed intervals of time on the water that were not between dives. We combined time engaged in diving and inter-dive intervals into a single behavioral category, representing active foraging. Thus, our final behavioral categories were: actively foraging (also referred to hereafter as “diving”), flying, at the colony, resting on sea ice, and resting on the water surface. For each T_b_ measurement, we determined which behavioral state the bird was in at that time by aligning time stamps from the T_b_ and behavioral (accelerometer) data in Microsoft Excel 16.16.27.

### Mercury contamination: sampling and analysis

We obtained small ∼0.2-0.5 ml blood samples from the brachial veins of focal individuals. Blood samples were centrifuged for 10 min. at 3500 rpm to separate plasma from red blood cells (RBCs), which were stored in 70% ethanol. RBCs were freeze dried for 48 hrs and homogenized prior to analysis for total Hg concentrations. Samples were analyzed in duplicate using an advanced Hg analyser spectrophotometer (Altec AMA 254) at the Institute Littoral Environnement et Sociétés, La Rochelle University (Bustamante et al. 2006). The standard deviation between duplicates was <10%. We used TORT-3 as a standardized reference material (CRM; Lobster Hepatopancreas Tort-3; NRC, Canada; [Hg] = 0.292 ± 0.022 µg g^−1^ dry weight (dw)) and performed a blank before initiating measurements on samples. The limit of detection for Hg and mean ± SD of Tort-3 measurements were 0.005 µg g^−1^ dw and 0.306 ± 0.004 µg g^−1^ dw, respectively.

### Statistical analysis

We conducted statistical analyses in R 3.6.1 (R Core Team, 2019). We used generalized additive mixed effect models (GAMMs) in R package mgcv (Wood 2011, 2017) to assess whether the mean T_b_ of little auks varied with behavioral activity classes, environmental conditions, or time of day. For this model, we used each observation of T_b_, while including appropriate random effects and correlation structure to account for non-independence of observations. Specifically, package mgcv allowed us to implement a correlation structure that accounts for temporal autocorrelation (corAR1 correlation structure implemented through package nlme; Pinheiro et al. 2019), to include individual ID and behavioral bought as random effects, and to incorporate a non-linear smooth term (cyclic cubic regression spline; specified as bs=cc) to account for potential variation in T_b_ with time of day. We included two-way interactions between behavioral class and: (1) ambient temperature, (2) relative humidity, (3) wind speed, (4) time spent engaged in the activity, and (5) Hg concentrations measured in the whole blood. These interactions test whether the relationship between T_b_ and behavior is contingent upon external conditions, the amount of time elapsed in a certain behavior (e.g. flying), and contamination level. We removed interactions with p-values > 0.050 from models, followed by elimination of main effects above the same threshold. We used the emtrends function in R package emmeans to test pairwise comparisons for interaction terms (Lenth 2019). Pairwise comparisons for mean differences in T_b_ between behavioral states were conducted using function emmeans within package emmeans. For this purpose, interactions were first removed from models to avoid complications with interpretation. In addition, to further explore the interaction that emerged between behavioral state and time spent engaged in the activity, we calculated change in T_b_ (deltas) for each behavioral bout as: ΔT_b_ = T_b,end_-T_b,start_; where T_b,end_ = T_b_ at the last time point recorded in that behavioral state and T_b,start_ = T_b_ at the first time point recorded. We then used a linear mixed effects model in nlme to compare ΔT_bs_ across behavioral states, and also included the length of the time interval in the model. We extracted and plotted predicted values from models using R function ggpredict within the ggeffects package (Lüdecke 2018).

In addition, we assessed whether between minute variation in T_b_ differed between behavioral states by calculating the absolute value of the difference between subsequent measurements of T_b_, and then constructing models with the same structure as described for mean T_b_. Values could not be calculated for time points at the beginning of the behavioral intervals, so these rows were dropped from the analysis.

## Results

### Effect of behavioral state on mean T_b_

The mean ± SD T_b_ of little auks across all observations was 41.02 ± 0.55 (range: 39.29-43.08). T_b_ varied significantly with behavioral state (*F*_*4*_ *=* 33.26; *P <* 0.001; Table 1a). Without interactions in the model, T_b_ was highest when birds were at the colony, followed by flying, and was lowest when birds were resting on the sea ice. T_b_ while birds were diving versus resting on the water surface did not significantly differ, and was intermediate to T_b_ while flying and on the ice. Pair-wise comparisons indicated significant differences in T_b_ in these states, with the exception of between diving and resting on the water (Table 1b). Figure 1 shows a representative trace of T_b_ variation through time for one focal individual. See Fig. S1-S7 for equivalent traces for the other 7 birds.

**Table 1.** Differences in mean T_b_ of little auks in the five behavioral states: (a) estimated marginal (EM) means from the best-fitting GAMM with interactions removed (df = 16391), (b) pairwise contrasts between behavioral states.

| (a) Behavioral state | EM mean ± SE [CI] | | |
| --- | --- | --- | --- |
| Diving (D) | 41.15 ± 0.046 [41.06, 41.24] | | |
| Flying (F) | 41.27 ± 0.043 [41.18, 41.35] | | |
| Colony (C) | 41.64 ± 0.050 [41.54, 41.74] | | |
| Ice (I) | 40.96 ± 0.056 [40.85, 41.07] | | |
| Water (W) | 41.14 ± 0.051 [41.04, 41.24] | | |
| (b) Pairwise |  |  |  |
| contrast | Estimate ± SE | *T* | *P* |
| D-F | -0.121 ± 0.038 | -3.192 | 0.012 |
| D-C | -0.495 ± 0.054 | -9.157 | <0.001 |
| D-I | 0.189 ± 0.050 | 3.752 | 0.002 |
| D-W | $0.008 \pm 0.043$ | 0.187 | 0.999 |
| F-C | $-0.374 \pm 0.053$ | -7.128 | <0.001 |
| F-I | $0.310 \pm 0.050$ | 6.156 | <0.001 |
| F-W | $0.129 \pm 0.043$ | 3.019 | 0.021 |
| C-I | $0.685 \pm 0.063$ | 10.92 | <0.001 |
| C-W | $0.503 \pm 0.058$ | 8.616 | <0.001 |
| I-W | $-0.181 \pm 0.053$ | -3.371 | 0.007 |

**Figure 1.**
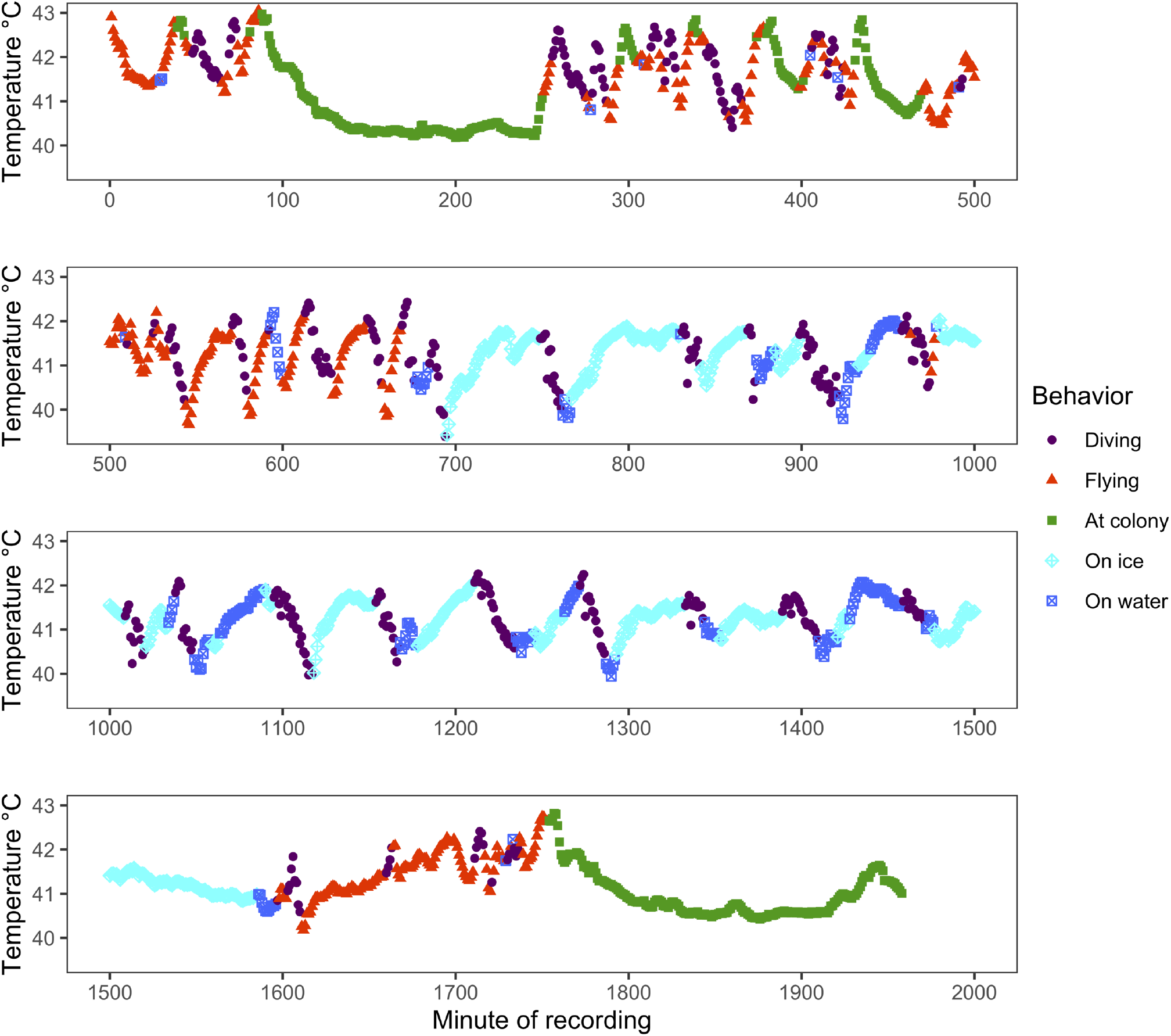
Variation in body temperature (T_b_) through time and color coded with respect to behavioral state for one individual little auk (LIAK20EG19) from the Ukaleqarteq, East Greenland, population. Note rebounds in T_b_ when on the sea ice following declines while diving in cold arctic waters. T_b_ can also be observed to increase with time during flight and decline with time at the colony.

### Effect of environmental conditions on mean T_b_

The best model predicting mean T_b_ included positive effects of ambient temperature (β ± SE = 0.008 ± 0.004, *T* = 2.012, *P* = 0.044; Fig. 2a) and relative humidity (β ± SE = 0.006 ± 0.001, *T* = 4.353, *P <* 0.001; Fig. 2b). The two-way interactions between ambient temperature, relative humidity and behavioral state were non-significant (*F*_4_ = 1.853, *P* = 0.116; *F*_4_ = 0.878, *P* = 0.476, respectively; Fig. 2a,b). There was a marginally significant interaction between wind speed and behavioral state in predicting T_b_ (*F*_4_ =2.302, *P* = 0.056; Fig. 2c). We proceeded to assess the meaning of this interaction by constructing models predicting the effect of wind speed within each behavioral state. The T_b_ of little auks increased with wind speed when birds were in flight, but did not vary with wind speed in the other behavioral states (Table 2a; Fig. 2c; see Table S1 for statistics for pairwise comparisons in the trends between behavioral states).

**Table 2.** Results of GAMMs predicting T_b_ within the behavioral states, showing estimated effects (Estimate ± SE, t, P) of wind speed (m/s) (a), and time interval (min) within the behavioral bought (b). Differences in superscript letters indicate significant differences between trends.

|  | (a) Wind speed | (b) Time interval |
| --- | --- | --- |
| Diving | -0.007 ± 0.008, -0.853, 0.393<sup>a</sup> | -0.024 ± 0.001, -15.15, <0.001<sup>a</sup> |
| Colony | 0.001 ± 0.005, 0.110, 0.913<sup>a,c</sup> | -0.004 ± 0.001, -5.807, <0.001<sup>b</sup> |
| Ice | 0.004 ± 0.004, 1.193, 0.233<sup>a,c</sup> | 0.014 ± 0.001, 12.36, <0.001<sup>c</sup> |
| Flying | 0.015 ± 0.007, 2.265, 0.0236<sup>a,c</sup> | 0.001 ± 0.0001, 1.580, 0.114<sup>c</sup> |
| Water | 0.001 ± 0.009, 0.148, 0.882<sup>b,c</sup> | 0.001 ± 0.003, 0.549, 0.583<sup>b,c</sup> |

**Figure 2.**
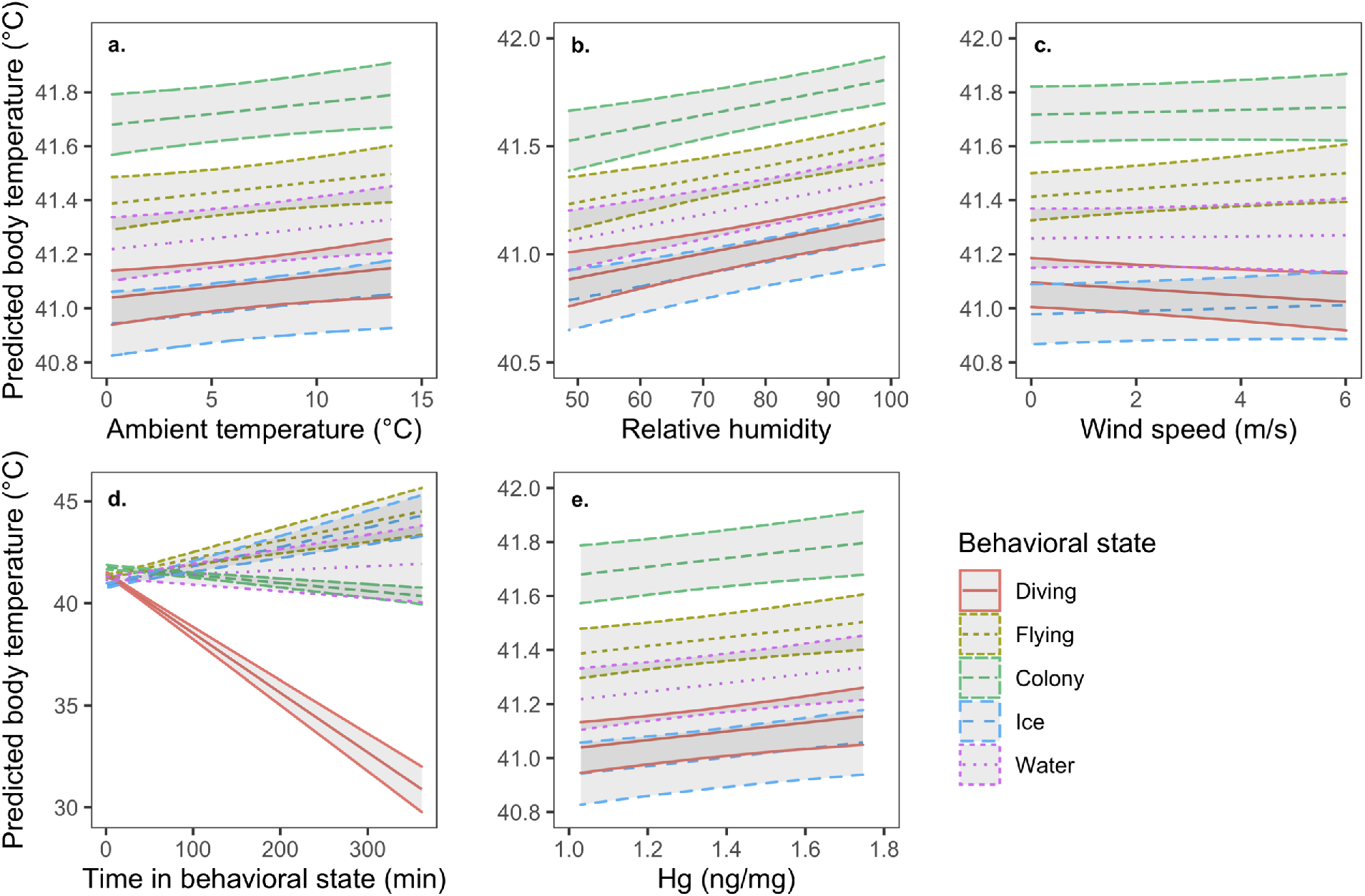
Relationships between the mean T_b_ of little auks and (a) ambient temperature, (b) relative humidity, (c) wind speed, (d) time within the behavioral state, and (e) mercury concentrations. The effect of wind speed and time in behavioral state on T_b_ varied with behavioral state, whereas the effects of the other variables were consistent across behavioral states.

### Interaction with time in the behavioral state in predicting mean T_b_ and ΔT_b_

In addition, the best model predicting the mean T_b_ of little auks included the interaction between behavioral state and time interval within the behavior state (*F*_4_ =49.23, *P* < 0.001; Fig. 1; Fig. 2d). We proceeded to assess the meaning of this interaction by constructing models predicting the effect of time interval within each behavioral state. The T_b_ of little auks decreased with the amount of time spent diving, or at the colony (Table 2b; Fig. 1; Fig. 2d). In contrast, T_b_ increased the longer birds spent on the sea ice and tended to increase during flight (Table 2b; Fig. 1; Fig. 2d). T_b_ did not consistently vary with time when birds were resting on the water surface (Table 2b; Fig. 1; Fig. 2d; see Table S2 for statistics for pairwise comparisons in trends between behavioral states).

Results regarding the ΔT_bs_ for the different behavioral states mostly aligned with the analysis above (Table S3). The ΔT_b_ for diving was negative, with the 95% CI not overlapping zero, and was significantly lower than all other delta T_bs_. In contrast, the ΔT_b_ for flying and resting on the sea ice were positive, with the CIs not overlapping zero, and were significantly higher than all other delta T_bs_, with the ΔT_b_ for sea ice also greater than that of flying. The ΔT_b_ for at the colony and on the water were negative, and positive, respectively, but did not significantly differ from each other or from zero (CIs overlapping zero) (Table S3),

### The effect of Hg on mean T_b_

The mean ± SE Hg concentration in the blood was 1.290 ± 0.031 µg g^−1^ dw [range: 1.030-1.746 µg g^−1^ dw], which assuming 79% blood moisture content is equivalent to 0.271 ± 0.007 µg g^−1^ ww [range: 0.216-0.367 µg g^−1^ ww], and falls within the range of low risk for toxicological effects (0.2–1.0 µg g^−1^ ww; Ackerman et al. 2016). Independent of behavioral state or weather conditions, the T_b_ of little auks was significantly higher in birds with higher Hg levels (β ± SE = 0.161 ± 0.061, *t =* 2.639, *P* = 0.008; Fig. 2e). Hg levels did not significantly interact with behavioral state (*F*_4_ = 1.811; *P* = 0.124), ambient temperature (*F*_1_ = 0.373, *P* = 0.542), relative humidity (*F*_1_ = 0.199, *P* = 0.655), or wind speed (*F*_1_ = 0.056, *P* = 0.812) to predict T_b_.

### Effect of time of day on mean T_b_

Mean T_b_ varied over the 24 hour period, with the best model including a highly non-linear (edf > 2) cyclic smooth spline term for the effect of time (edf = 5.544, *F*_8_ = 77.858, *P* < 0.001; Fig. 2). The spline term suggested a peak in T_b_ around mid-day (∼10:00-15:00) and the lowest values in the early morning (∼4:00 am) (Fig. 3).

**Figure 3.**
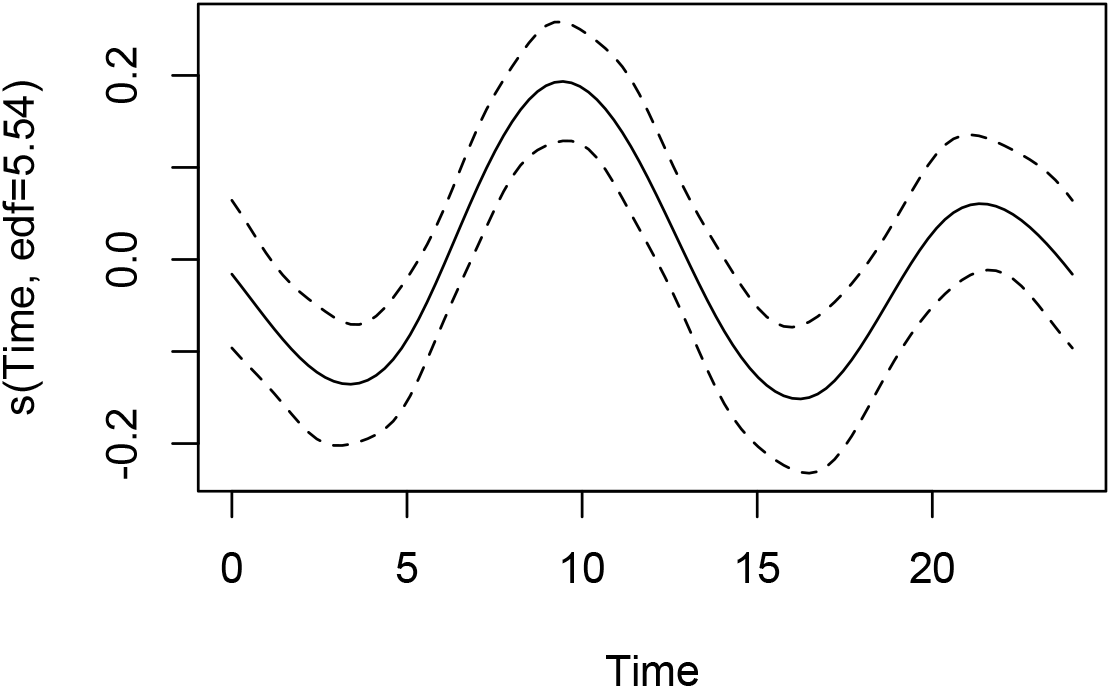
Residuals from the gamm model predicting mean T_b_ from time of day, with a smooth curve fitted. Dashed lines show 2-SE limits. The y-axis shows the partial effect of time on T_b_.

### Effect of behavioral state on between minute variation in T_b_

The mean ± SD of between minute variation in T_b_, as quantified by the absolute value of the difference between consecutive 1 minute readings, was 0.089 ± 0.106 (range: 0-1.46). Variation was highest when birds were foraging at sea, followed by flying, resting on the water surface, at the colony and on the sea ice (Table 3). There were significant differences in the between minute variation in T_b_ between behavioral states, with the exception of when birds were in flight and resting on the water surface (Table 3).

**Table 3.** Between minute variation (|T_b1_-T_b1+1_|) in T_b_ of little auks in the five behavioral states: (a) estimated marginal (EM) means from the best-fitting GAMM with interactions removed (*df* = 15127), (b) pairwise contrasts between behavioral states.

| (a) Behavioral |  |  |  |
| --- | --- | --- | --- |
| state | EM mean ± SE [CI] | | |
| Diving (D) | 0.156 ± 0.003 [0.149, 0.163] | | |
| Flying (F) | 0.118 ± 0.004 [0.110, 0.126] | | |
| Colony (C) | 0.091 ± 0.006 [0.079, 0.103] | | |
| Ice (I) | 0.063 ± 0.006 [0.052, 0.074] | | |
| Water (W) | 0.113 ± 0.005 [0.103, 0.124] | | |
| (b) Pairwise |  |  |  |
| contrast | Estimate ± SE | *T* | *P* |
| D-F | 0.038 ± 0.055 | 6.890 | <0.001 |
| D-C | 0.065 ± 0.007 | 9.023 | <0.001 |
| D-I | 0.093 ± 0.006 | 14.18 | <0.001 |
| D-W | 0.043 ± 0.006 | 6.663 | <0.001 |
| F-C | 0.027 ± 0.008 | 3.622 | 0.003 |
| F-I | 0.055 ± 0.007 | 8.053 | <0.001 |
| F-W | 0.005 ± 0.006 | 0.694 | 0.958 |
| C-I | 0.028 ± 0.008 | 3.381 | 0.007 |
| C-W | -0.023 ± 0.008 | -2.759 | 0.046 |
| I-W | -0.051 ± 0.008 | -6.683 | <0.001 |

### Effect of environmental conditions on between minute variation in T_b_

The best model describing between minute variation in T_b_ included a positive effect of ambient temperature (± SE = 0.003 ± 0.001, *T* = 4.212, *P* < 0.001; Fig. 4a). There was also a significant interaction between wind speed and behavioral state in predicting variation in T_b_ (*F*_4_ = 4.99, *P* < 0.001; Fig. 4b). Again, we assessed the meaning of this interaction by constructing models predicting the effect of wind speed within each behavioral state. Variation in T_b_ increased with wind speed when birds were diving and flying. In contrast, wind speed was not significantly related to variation in T_b_ when birds were at the colony, on the sea ice, or resting on the water surface, and the coefficient estimate within these behavioral states was negative (Table 4a; see Table S4 for statistics for pairwise comparisons in the trends between behavioral states). Relative humidity was not related to variation in T_b_ (± SE = 0.0001 ± 0.0003, *T* = 0.351, *P* = 0.723), and the interactions between ambient temperature, relative humidity, and behavioral state were non-significant (*F*_4_ = 0.096, *P* = 0.983; *F*_4_ = 0.052, *P* = 0.995; respectively).

**Table 4.** Results of GAMMs predicting variation (|T_b1_-T_b1+1_|) in T_b_ within the behavioral states, showing estimated effects (Estimate ± SE, t, P) of wind speed (m/s) (a), and time interval (min) within the behavioral bought (b). Differences in superscript letters indicate significant differences between trends.

|  | (a) Wind speed | (b) Time interval |
| --- | --- | --- |
| Diving | $0.009 \pm 0.003$, 3.054, 0.002<sup>a</sup> | $-0.003 \pm 0.0002$, -10.99, <0.001<sup>a</sup> |
| Colony | $-0.003 \pm 0.002$, -1.690, 0.091<sup>b</sup> | $-0.0001 \pm 0.0003$, -2.586, 0.009<sup>b</sup> |
| Ice | $-0.001 \pm 0.002$, -0.533, 0.594<sup>b</sup> | $-0.0006 \pm 0.0001$, -9.966, <0.001<sup>c</sup> |
| Flying | $0.007 \pm 0.003$, 2.585, 0.010<sup>a,b</sup> | $-0.002 \pm 0.0002$, -8.580, <0.001<sup>d</sup> |
| Water | $-0.006 \pm 0.004$, -1.608, 0.108<sup>b</sup> | $-0.003 \pm 0.0004$, -8.557, <0.001<sup>a</sup> |

**Figure 4.**
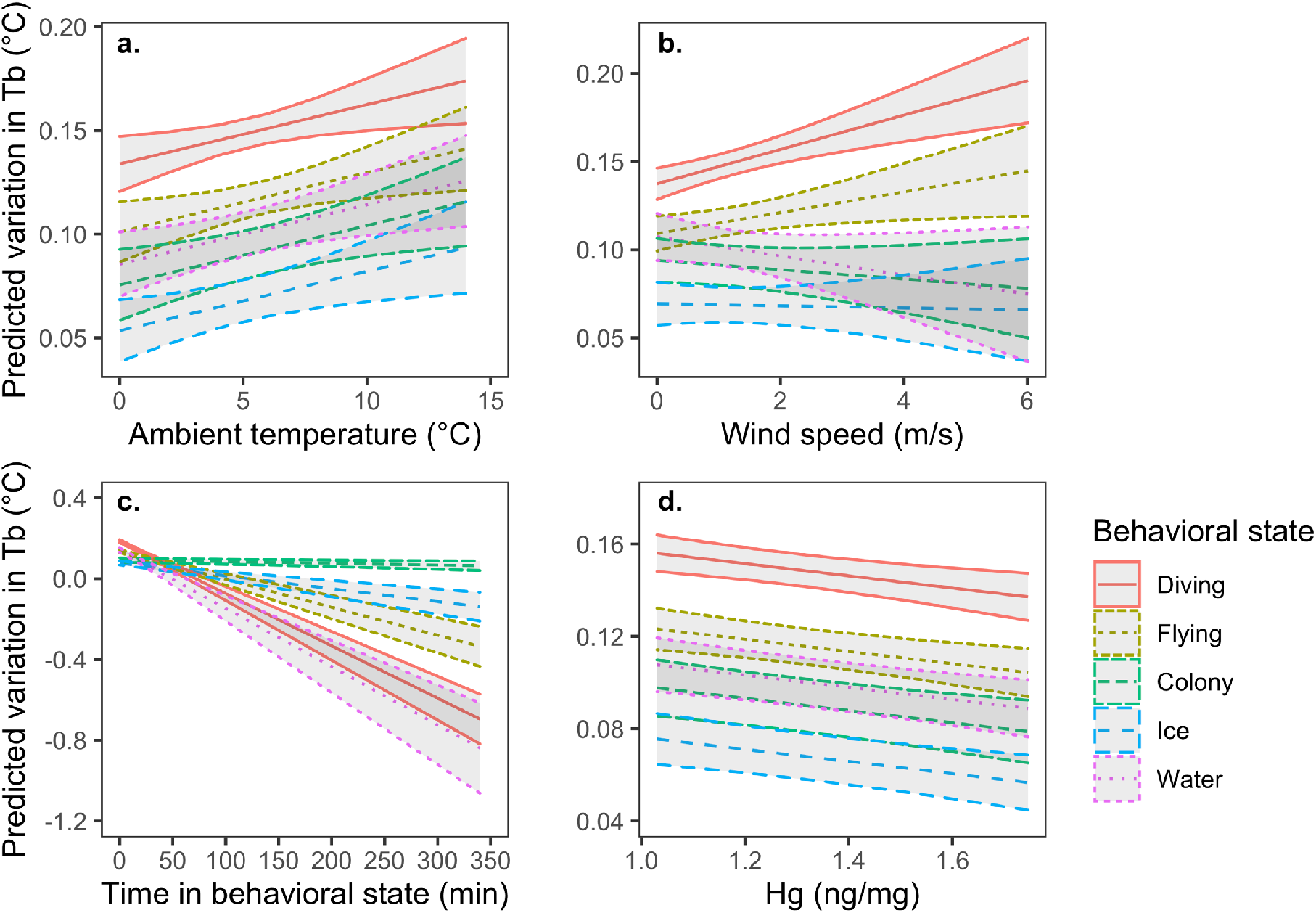
Relationships between the between minute variation in T_b_ of little auks and (a) ambient temperature, (b) wind speed, (c) time within the behavioral state, and (e) mercury concentrations. The effect of wind speed and time in behavioral state on T_b_ varied with behavioral state, whereas the effects of the other variables were consistent across behavioral states.

### Interaction with time in the behavioral state in predicting between minute variation in T_b_

As for mean T_b_, there was an interaction between time interval within the behavioral state and behavioral state in predicting between minute variation in T_b_ (*F*_4_ = 75.77, *P* < 0.001; Fig. 4c). Models constructed within the behavioral states indicated that between minute variation in T_b_ decreased with time in the behavioral state for all behaviors. However, this decrease was the steepest, and relatively equal in magnitude, when birds were resting in the water or engaged in diving bouts. The next steepest decrease was observed when birds were in flight, followed by when birds were on the sea ice, and the decrease was lowest when birds were at the colony (Table 4b; see Table S5 for statistics for pairwise comparisons in the trends between behaviors states).

### The effect of mercury on between minute variation in T_b_

Mercury concentrations in the blood did not significantly interact with behavioral state (*F*_4_ = 0.842; *P* = 0.498), ambient temperature (*F*_1_ = 0.373, *P* = 0.542), relative humidity (*F*_1_ = 3.154, *P* = 0.076), or wind speed (*F*_1_ = 0.214, *P* = 0.643) to predict between minute variation in T_b_. However, independent of behavioral state or weather conditions, between minute variation in the T_b_ of little auks was significantly lower in birds with higher Hg levels (± SE = −0.026 ± 0.008, *t = -*3.193, *P* = 0.001; Fig. 4d).

### Effect of time of day on between minute variation in T_b_

Between minute variation in T_b_ did not vary with time of day (edf = 0.949, *F*_8_ = 0.226, *P* = 0.127).

## Discussion

Although T_b_ in little auks was relatively tightly regulated around a mean of 41.02 °C, we observed low, but significant, variation according to behavioral state and weather conditions. Such plasticity may be critical to maintaining energy balance across contexts and may buffer species against negative fitness effects in the context of global change. Nevertheless, the potential for plasticity to prevent energetic costs is not infinite, and our study suggested potential thermoregulatory challenges under global climate change scenarios. In particular, T_b_ rebounded on sea ice following declines during diving episodes in frigid Artic waters, suggesting that loss of this resting substrate may elevate thermoregulatory costs, negatively affecting energy balance, body condition, and/or fitness. In addition, we tested whether exposure of little auks to Hg interacted with environmental conditions to affect T_b_. Hg is an important chemical contaminant in the Arctic, and has the potential to disrupt thermoregulatory processes, for instance through endocrine disruption, or by imposing detoxification challenges. No interactive effects between Hg exposure and environmental conditions were detected. However, a concerning contingency is that the higher, less variable, T_b_ observed in more contaminated birds could limit plasticity and/or pose energetic challenges under future scenarios of global change.

The most exciting finding of our study was that T_b_ rebounded when birds were resting on sea ice, following declines while foraging in cold Arctic waters. Specifically, average T_b_ on the sea ice was actually lower than in any other behavioral state, which we believe reflects the fact that birds exit the water when their T_b_ falls below a threshold. While resting on sea ice, birds increased their T_b_ by an average of ∼0.33°C, meaning that, on average, they approximately recovered the amount of T_b_ lost while foraging, that is, ∼0.31°C. In contrast, while resting on the water surface, T_b_ remained relatively unchanged and did not differ significantly from when birds were foraging. Furthermore, the lowest variation in T_b_ was observed when birds were resting on sea ice, whereas the highest was observed during bouts of diving behavior, supporting the hypothesis that sea ice plays an important role in allowing birds to restore and maintain normiothermic temperatures after thermally challenging foraging bouts.

In the context of climate change, loss of sea ice may have significant energetic and thermoregulatory implications, as birds are forced to instead rest on the water surface. Birds face increased thermoregulatory challenges when in the water, which has much higher (∼25 ×) thermal conductivity than air (Grémillet et al. 2015). In auks, compression of air space in the feathers while diving significantly reduces insulative properties, further facilitating heat exchange with the environment (Oswald and Arnold 2012). Thus, birds resting on the water likely elevate their metabolic rate, even considering lower T_b_, which, *in lieu* of compensatory changes in behavior or physiology, could elevate daily energy expenditure and threaten to result in negative energy balance (Lovvorn et al. 2009). In addition, if T_b_ does not increase while resting on the water, but does when birds are at rest on sea ice, this could force birds to return to the colony sooner, limiting time for energy acquisition. Loss of sea ice as a substrate for resting, foraging, and movement has demonstrated effects on energy balance and population dynamics in many sea ice-dependent species (Post et al. 2013; Laidre et al. 2020; Pagano and Williams 2021). For instance, polar bears (*Ursus maritimus*) and narwhal (*Monodon monoceros*), which both have foraging ecologies tightly with sea ice, show substantial elevations in locomotory costs (3-4 times greater) in association with sea ice declines (Pagano and Williams 2021). However, effects of sea ice loss on thermoregulatory dynamics have been under-explored.

A second major result of our study was demonstrating sensitivity of little auks’ T_b_ to external environmental conditions, which may have implications under global change scenarios. Mean T_b_ increased with ambient temperature and relative humidity, with these relationships consistent across behavioral states. As ambient temperature and relative humidity rise, the capacity for evaporative heat dissipation decreases, which may result in increases in T_b_ or elevated thermoregulatory costs to maintain optimal T_b_ (Dawson 182; Gerson et al. 2014). In contrast, T_b_ increased with wind speed only when birds were in flight, demonstrating that high winds increase the thermal challenges of flying, but have little effect on thermodynamics during other activities. With respect to the environmental-sensitivity of between minute variation in T_b_, we observed an increase with ambient temperature, independent of behavioral state, suggesting that these cold-adapted birds face increasing challenges maintaining a stable T_b_ at higher temperatures. In addition, as for mean T_b_, we observed an interaction between wind speed and behavioral state, with between minute variation in T_b_ increasing with wind speed when birds were diving and flying, perhaps reflecting increased energetic and thermal challenges associated with activity during high winds. In contrast, wind speed was not strongly related to variation in T_b_ during the other behavioral states, which entail lower activity levels. In the context of climate change, these results suggest that alterations in ambient temperature may have implications for T_b_ regulation that are relatively independent of behavioral state, whereas changes in wind patterns may have especially high costs during active periods, especially during flight.

A third important finding of our study was that mean levels, and between minute variation, in T_b_ varied with concentrations of Hg in the blood. Independent of behavioral state, T_b_ increased with Hg levels. The elevation in T_b_ observed in birds with higher Hg levels could reflect an elevation in basal metabolic rate (BMR) as a result of detoxification costs, such as those associated with depuration (Gerson et al. 2019b). The association between Hg exposure and BMR remains poorly explored and equivocal in wildlife (Chastel et al. 2022), and in the only study to invest the relationship between Hg contamination and BMR in Arctic birds, Hg was unassociated with BMR (Blévin et al. 2017). However, in a laboratory study on zebra finch (*Taeniopygia guttata*), exposure to environmentally-relevant levels of MeHg were associated with elevated BMR (Gerson et al. 2019b). To our knowledge, there is no previous study documenting a link between Hg concentration and T_b_ in free-ranging animals. However, contrary to our results, studies in laboratory animals have demonstrated hypothermic responses to Hg exposure, for instance, in the mouse (*Mus musculus*) (Gordon et al. 1990). Hypothermic responses to chemical contamination are hypothesized to reduce the toxicity of the contaminant in the body (Leon 2008), but may not be relevant at the relatively low Hg levels in our little auk population. In addition, since our results our correlational, the association between Hg contamination levels and T_b_ could be indirect, for instance, reflecting a higher rate of Hg accumulation in individuals with intrinsic differences in BMR and feeding rate.

In contrast to for mean T_b,_ we observed a negative relationship between Hg levels and minute-by-minute variation in T_b_, which is inconsistent with the hypothesis that contaminated bird could have more difficulty maintaining a stable T_b_. Moreover, Hg levels did not interact with environmental conditions to affect T_b_, thus providing no evidence that exposure to this contaminant accentuates thermoregulatory responses to environmental change. However, absence of such interactive effects could reflect limitations to the range of Hg levels and environmental conditions spanned by our study. There is need for more research to examine the potential for interactive effects at higher contamination levels, and across steeper environmental gradients.

In addition to these central findings, our results suggested that the colony, as well as sea ice, serves as a thermal refuge for little auks, and indeed, may be selected for thermo-protective properties. The highest average levels of T_b_ were observed while birds were at the colony, even higher than when birds were in flight. Furthermore, the second lowest between minute variation in T_b_ occurred at the colony, which is consistent with the colony providing stable thermal conditions for birds. Higher average T_b_ at the colony than when in flight, which opposes our predictions, may be due to a combination of factors. First, little auks occupy a relatively cold environment in which thermoregulatory substitution may occur. That is, heat generated in flight may offset thermoregulatory costs, reducing changes in T_b_ with activity (Careau and Garland 2012). Second, birds at the colony may be buffered from the effects of movement and wind exposure that disrupt the boundary layer and increase heat flux from the body. However, T_b_ did decrease slightly the longer birds spent at the colony, which may reflect an adaptive downregulation in resting birds or decreases in T_b_ following commuting flights between foraging sites and the colony.

As expected, we observed that T_b_ was higher when little auks were flying than when they were diving or resting on the water. Furthermore, although mean T_b_ in flight was lower than at the colony, T_b_ in flight increased, suggesting that heat generated during flight does increase T_b_. For little auks, this pattern is expected due to their unique morphology. Little auks can be described as bullets with wings. They flap or fall. Their morphology and flight mode are characterized by high wing loadings and rapid wing beats, which translates into high flight costs (estimated at ∼7.24 x basal metabolic rate; Ste-Marie et al. 2022).

Also in-line with expectations, and discussed to some extent above, we observed that the T_b_ declined while little auks were foraging in cold waters. This decline in T_b_ may facilitate aerobic capacity during diving and limit heat loss, but may also reflect unavoidable declines linked to submergence in cold water with a high thermal conductance (Kooyman and Ponganis 1998; Favilla and Costa 2020; Williams and Ponganis 2021). In addition, declines in T_b_ while foraging may be magnified by ingestion of cold prey items. Indeed, ingestion-linked declines in T_b_ have been used to identify feeding events by past studies (Wilson et al. 1995). Our T_b_ data was logged at 1-min intervals, and thus lacked the resolution necessary to identify changes in T_b_ associated with independent feeding events. In another species of Alcid, the Brünnich’s guillemots (*Uria lomvia*), T_b_ was observed to decline over the course of sequential diving bouts, as we see in the little auks. However, this decline occurred during periods resting on the water between dives, rather than during dives themselves. In the guillemots, T_b_ increased during dives, while the temperature of the periphery declined. This pattern contrasts to the declines in T_b_ during diving which have been observed in some species of penguins (Bevan et al. 2002; Green et al. 2003; Williams and Ponganis 2021), and was interpreted as reflecting a combination of peripheral vasoconstriction and high wing beat frequency that generates heat (Niizuma et al. 2007). Our data lack the resolution to effectively separate dives from inter-dive intervals. Thus, a similar dynamic could also be occurring in our birds.

Finally, the best model predicting mean T_b_ included a non-linear effect of time, with the highest values occurring around mid-day and the lowest values in the early morning. Despite the fact that little auks in our population breed under 24-hrs of daylight, this diurnal variation in T_b_ may reflect timing of maximum solar radiation exposure, a diel pattern of activity levels, and/or underlying circadian rhythmicity in T_b_ independent of activity levels. A past study on little auks found a regular rhythm of population attendance at the population level, likely associated with period of lower predation pressure, which provides some foundation for expecting that T_b_ could also display a diel pattern of variation. However, this same study found little circadian rhythm in activity patterns of individual little auks (Wojczulanis-Jakubas et al. 2020), In contrast to mean T_b_, between minute variation in T_b_ did not show a pattern with time of day.

## Conclusions

Our results demonstrate that the T_b_ of little auks is modulated according to both behavioral state and environmental conditions, which likely aids animals in maximizing energy balance while performing essential behaviors in dynamic environments. Although this plasticity is predicted to facilitate energy balance in the face of climate change, the dynamic nature of T_b_ regulation also suggests that changing environmental conditions may significantly alter energy balance, or the behavioral and energetic strategies that must be adopted to achieve energetic homeostasis. Most notably, our data suggests that little auks use sea ice as a thermal refuge, resting on this substrate to allow T_b_ to rebound after submersion in cold water. If sea ice is lost due to warming temperature, thermoregulatory costs are forecast to increase substantially as birds are forced to instead rest on the water surface. Furthermore, the elevated and less variable T_b_ of little auks with high Hg concentrations is of potential concern. Higher T_b_ in contaminated birds could elevate energetic costs or limit plasticity, further challenging scope for maintaining energy balance under scenarios of environmental change.

## Data availability

Data will be made available via the Zenodo community of European Commission Funded Research (OpenAIRE) (doi: 10.5281/zenodo.7220883).

## Acknowledgement

We thank Valère Marsaudon for his help with data collection in East Greenland, and members of Nanu travel for logistical support. We are grateful to Clément Bertin for aid in extracting sea ice coverage data. MLG and ALG are supported by the European Union’s Horizon 2020 programme (Marie Skłodowska-Curie grants 101025549, 896866). We acknowledge long-term support from the French Polar Institute (IPEV), through the ADACLIM program (388) administered by JF and DG. This work contributes to research projects ARCTIC-STRESSORS and ILETOP funded by the French National Research Agency (ANR-20-CE34-0006, ANR-16-CE34-0005), the international initiative ARCTOX (arctox.cnrs.fr) and the Excellence Chair ECOMM funded by the Region Nouvelle Aquitaine.

## Authors’ contributions

MLG, ASG and JF conceived the study. MLG, ASG, DG and JF obtained funding for fieldwork and laboratory analyses. MSG, ALG, SG and JF collected the data. MLG, ASG and AS analysed the data. MLG and ASG wrote the first draft of the manuscript. All authors read and commented on the manuscript.

