## Supplementary Material for "Body temperature rebounds on sea ice and is elevated by mercury contamination in a keystone predator"

^e^Institut Universitaire de France (IUF), 1 rue Descartes 75005 Paris, France

**Caption**: In this manuscript, we explore how variation in the body temperature (T_b_) of a keystone Arctic seabird, the little auk (*Alle alle*), is related to five behavioral states (flying, foraging (diving), at the colony, on sea ice, resting on the water), as assessed through 3D accelerometry, and environmental conditions. In addition, we evaluate whether mercury (Hg) contamination levels alter patterns of plasticity in T_b_, which may aid birds in coping with climate change-related alterations in environmental conditions, while maintaining behaviors critical to fitness. In this supplement, we present results of statistical analyses that are too extensive for inclusion in the main text (Table S1-S5). In addition, we include graphs (Fig. S1-S7) showing variation in T_b_ through time, and color-coded with respect to behavioral state) for 7 of our 8 focal birds (the graph for the 8^th^ individual is included as a representative in the main manuscript).

**Table S1.** Pairwise comparisons for the slopes of the relationship between mean body temperature and wind speed (m/s) in the five behavioral states. Extracted from the best-fitting GAMM (*df*=16381).

| (a) Behavioral state | EM trend ± SE [95% CI] | | |
| --- | --- | --- | --- |
| Diving (D) | -0.012 ± 0.006 [-0.024, <0.001] | | |
| Flying (F) | 0.015 ± 0.007 [0.001, 0.028] | | |
| Colony (C) | 0.005 ± 0.007 [-0.009, 0.018] | | |
| Ice (I) | 0.006 ± 0.007 [-0.007, 0.019] | | |
| Water (W) | 0.002 ± 0.008 [-0.015, 0.019] | | |
| (b) Pairwise contrast | Estimate ± SE | *T* | *P* |
| D-F | -0.026 ± 0.009 | -2.391 | 0.028 |
| D-C | -0.016 ± 0.009 | -1.767 | 0.393 |
| D-I | -0.017 ± 0.009 | -1.955 | 0.288 |
| D-W | -0.014 ± 0.010 | -1.298 | 0.693 |
| F-C | 0.010 ± 0.009 | 1.029 | 0.842 |
| F-I | 0.009 ± 0.009 | 0.940 | 0.881 |
| F-W | 0.013 ± 0.011 | 1.158 | 0.775 |
| C-I | -0.001 ± 0.009 | -0.118 | 1.000 |
| C-W | 0.002 ± 0.011 | 0.241 | 0.999 |
| I-W | 0.004 ± 0.011 | 0.351 | 0.997 |

**Table S2.** Pairwise comparisons for the slopes of the relationship between mean body temperature and time in the behavioral bought (min) in the five behavioral states. Extracted from the best-fitting GAMM (*df*=16381).

| (a) Behavioral state | EM trend ± SE [95% CI] | | |
| --- | --- | --- | --- |
| Diving (D) | -0.029 ± 0.001 [-0.032, -0.026] | | |
| Flying (F) | 0.008 ± 0.002 [0.006, 0.012] | | |
| Colony (C) | -0.003 ± 0.001 [-0.005, -0.002] | | |
| Ice (I) | 0.009 ± 0.001 [0.007, 0.012] | | |
| Water (W) | 0.002 ± 0.003 [-0.003, 0.007] | | |
| (b) Pairwise contrast | Estimate ± SE | *T* | *P* |
| D-F | -0.038 ± 0.002 | -16.55 | <0.001 |
| D-C | -0.025 ± 0.001 | -14.72 | <0.001 |
| D-I | -0.038 ± 0.002 | -17.72 | <0.001 |
| D-W | -0.031 ± 0.003 | -9.992 | <0.001 |
| F-C | 0.013 ± 0.001 | 7.242 | <0.001 |
| F-I | -0.001 ± 0.002 | -0.305 | 0.998 |
| F-W | 0.007 ± 0.003 | 2.203 | 0.179 |
| C-I | -0.013 ± 0.001 | -8.364 | <0.001 |
| C-W | -0.005 ± 0.003 | -2.117 | 0.213 |
| I-W | 0.007 ± 0.003 | 2.483 | 0.094 |

**Table S3**. Differences in mean ΔT_b_ of little auks in the five behavioral states: (a) estimated marginal (EM) means from the LMM (df = 1115), (b) pairwise contrasts between behavioral states.

| (a) Behavioral state | EM mean ± SE [CI] | | |
| --- | --- | --- | --- |
| Diving (D) | -0.312 ± 0.026 [-0.374, -0.251] | | |
| Flying (F) | 0.146 ± 0.029 [0.077, 0.215] | | |
| Colony (C) | -0.074 ± 0.044 [-0.179, 0.031] | | |
| Ice (I) | 0.335 ± 0.045 [0.228, 0.443] | | |
| Water (W) | 0.0002 ± 0.038 [-0.089, 0.089] | | |
| (b) Pairwise contrast | Estimate ± SE | *T* | *P* |
| D-F | -0.459 ± 0.045 | -10.29 | <0.001 |
| D-C | -0.238 ± 0.050 | -4.747 | <0.001 |
| D-I | -0.648 ± 0.055 | -11.70 | <0.001 |
| D-W | -0.313 ± 0.049 | -6.383 | <0.001 |
| F-C | 0.221 ± 0.058 | 3.737 | 0.002 |
| F-I | -0.189 ± 0.055 | -3.463 | 0.005 |
| F-W | 0.146 ± 0.049 | 2.991 | 0.024 |
| C-I | -0.409 ± 0.063 | -6.546 | <0.001 |
| C-W | -0.074 ± 0.058 | -1.273 | 0.708 |
| I-W | 0.335 ± 0.064 | 5.212 | <0.001 |

**Table S4.** Pairwise comparisons for the slopes of the relationship between variation in body temperature and wind speed (m/s) in the five behavioral states. Extracted from the best-fitting GAMM (*df*=15119).

| (a) Behavioral state | EM trend ± SE [95% CI] | | |
| --- | --- | --- | --- |
| Diving (D) | 0.009 ± 0.002 [0.005, 0.014] | | |
| Flying (F) | 0.006 ± 0.003 [0.001, 0.011] | | |
| Colony (C) | -0.003 ± 0.003 [-0.008, 0.002] | | |
| Ice (I) | -0.0006 ± 0.003 [-0.006, 0.005] | | |
| Water (W) | -0.005 ± 0.003 [-0.013, 0.002] | | |
| (b) Pairwise contrast | Estimate ± SE | *T* | *P* |
| D-F | 0.004 ± 0.003 | 1.083 | 0.815 |
| D-C | 0.012 ± 0.004 | 3.503 | 0.004 |
| D-I | 0.010 ± 0.004 | 2.718 | 0.051 |
| D-W | 0.015 ± 0.004 | 3.339 | 0.008 |
| F-C | 0.009 ± 0.003 | 2.343 | 0.131 |
| F-I | 0.006 ± 0.004 | 1.654 | 0.463 |
| F-W | 0.011 ± 0.005 | 2.439 | 0.105 |
| C-I | -0.002 ± 0.004 | -0.548 | 0.982 |
| C-W | 0.002 ± 0.004 | 0.579 | 0.978 |
| I-W | 0.004 ± 0.004 | 1.001 | 0.855 |

**Table S5.** Pairwise comparisons for the slopes of the relationship between variation in body temperature and time in the behavioral bought (min) in the five behavioral states. Extracted from the best-fitting GAMM (*df*=15119).

| (a) Behavioral state | EM trend ± SE [95% CI] | | |
| --- | --- | --- | --- |
| Diving (D) | -0.003 ± 0.0002 [-0.003, -0.002] | | |
| Flying (F) | -0.001 ± 0.0002 [-0.002, -0.001] | | |
| Colony (C) | -0.0001 ± 0.00008 [-0.0001, -0.00002] | | |
| Ice (I) | -0.0006 ± 0.0001 [-0.0009, -0.0004] | | |
| Water (W) | -0.003 ± 0.0003 [-0.004, -0.002] | | |
| (b) Pairwise contrast | Estimate ± SE | *T* | *P* |
| D-F | -0.001 ± 0.0002 | -4.952 | <0.001 |
| D-C | -0.003 ± 0.0002 | -13.01 | <0.001 |
| D-I | -0.002 ± 0.0002 | -8.882 | <0.001 |
| D-W | 0.0003 ± 0.0004 | 0.739 | 0.947 |
| F-C | -0.001 ± 0.0002 | -8.355 | <0.001 |
| F-I | -0.0007 ± 0.0002 | -3.968 | <0.001 |
| F-W | 0.001 ± 0.0003 | 3.982 | <0.001 |
| C-I | 0.0005 ± 0.0001 | 4.785 | <0.001 |
| C-W | 0.002 ± 0.0003 | 8.114 | <0.001 |
| I-W | 0.002 ± 0.0004 | 6.220 | <0.001 |

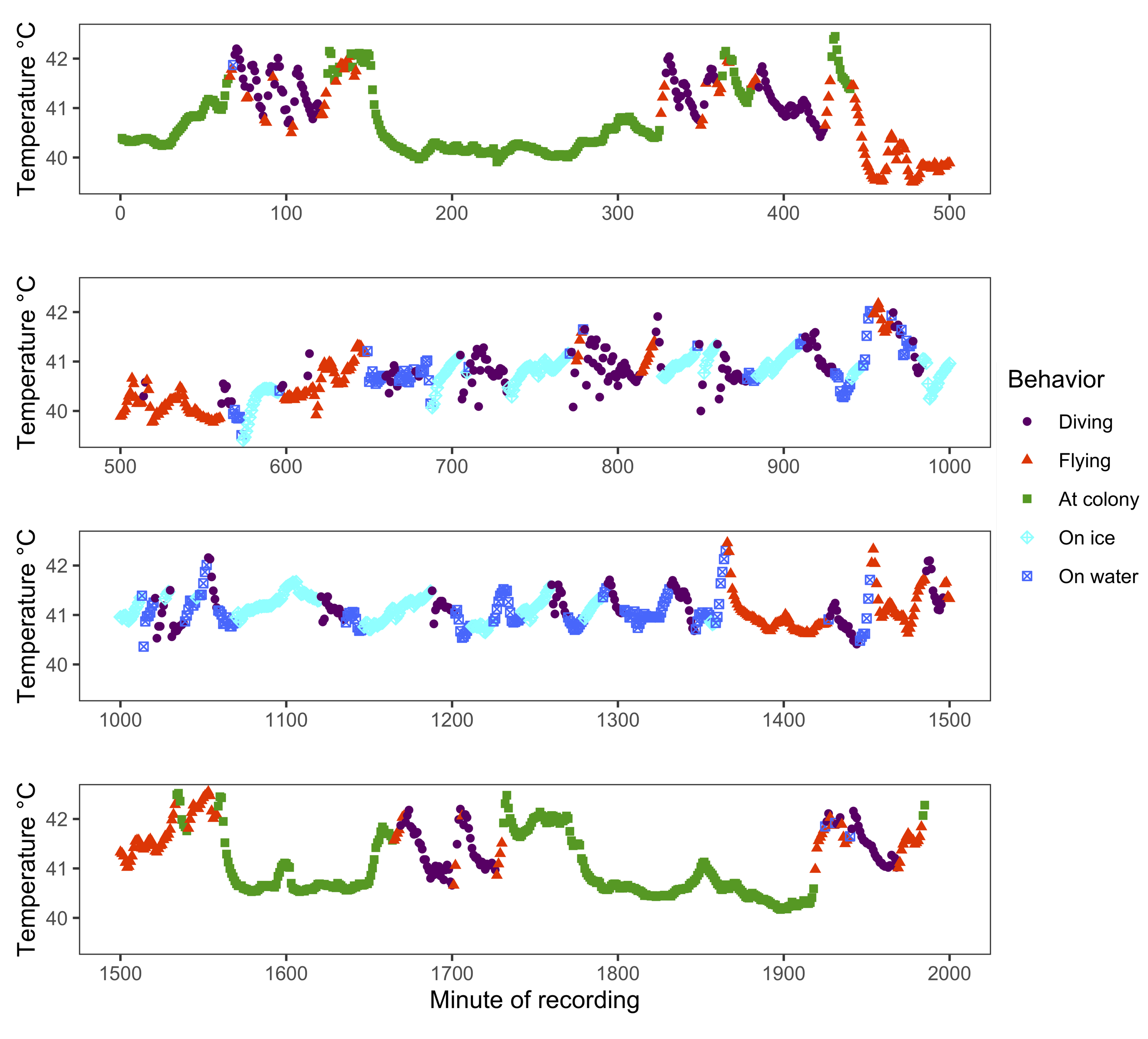

Fig S1. Variation in body temperature (T_b_**)** through time and color coded with respect to behavioral state for one individual little auk (LIAK20EG09) from the Ukaleqarteq, East Greenland, population. In this individual, decreases in T_b_ while diving, followed by rebounds in T_b_ on the sea ice are clearly notable, and T_b_ also appears to decrease with time at the colony. Although the overall pattern is for T_b_ to tend to increase with time in flight, in this individual, there are also some instances wherein T_b_ appears to decrease while flying. Across individuals, such instances appear most common when birds are departing the colony (e.g. around 450 min), and could reflect exposure to colder air temperatures and wind over the sea relative to over land.

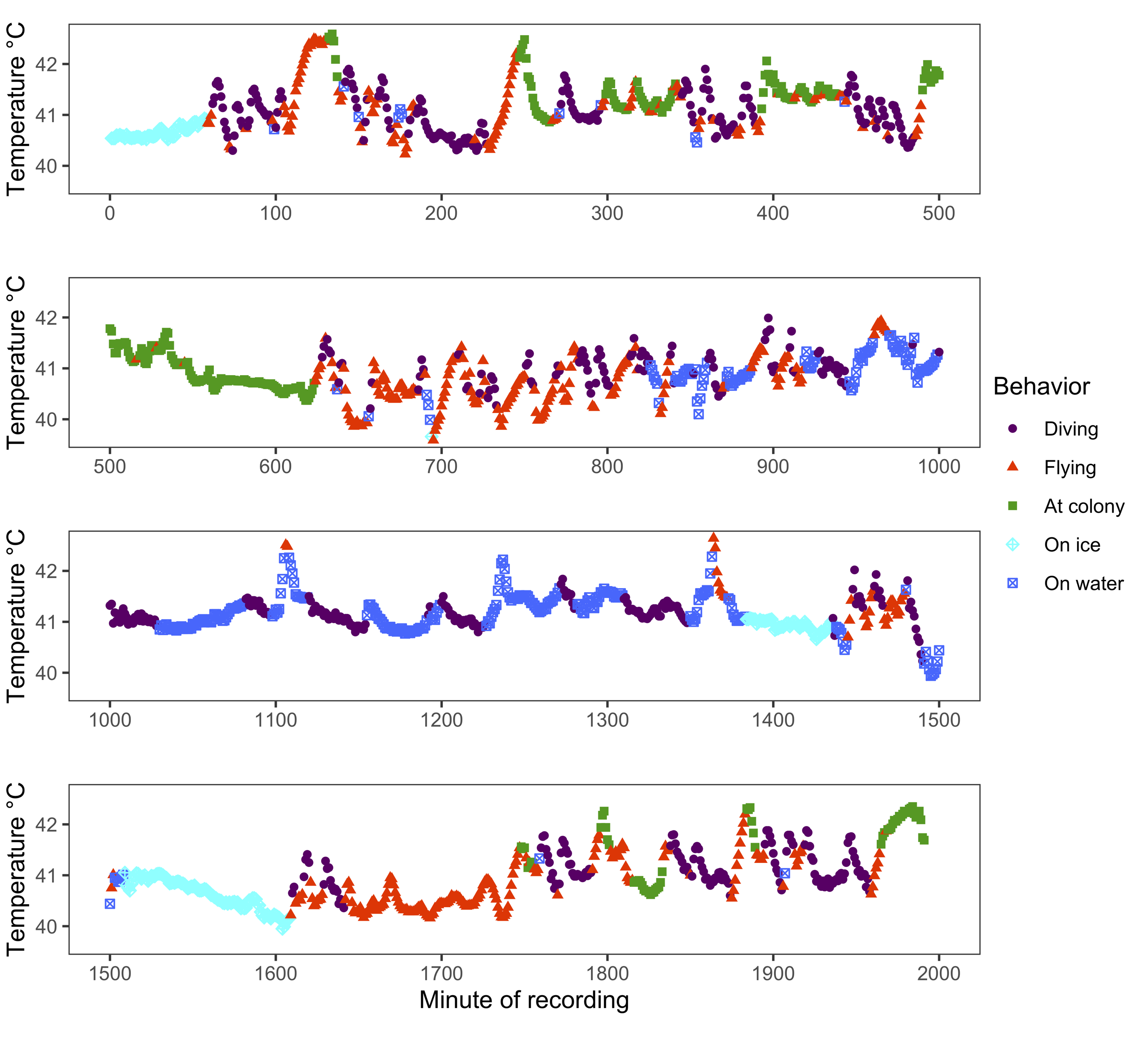

Fig. S2. Variation in body temperature (T_b_**)** through time and color coded with respect to behavioral state for one individual little auk (LIAK20EG12) from the Ukaleqarteq, East Greenland, population. In this individual, the time spent on sea ice does not immediately follow diving bouts, so the rebound in T_b_ on sea ice, which observed in other individuals, is not apparent. Notably, rather than spending time on sea ice in between diving, the individual spent more time on water, and it can be observed that T_b_ does recover as well while the bird is resting on the water, relative to on sea ice. T_b_ can be observed to generally increase during flight, and decrease while at the colony.

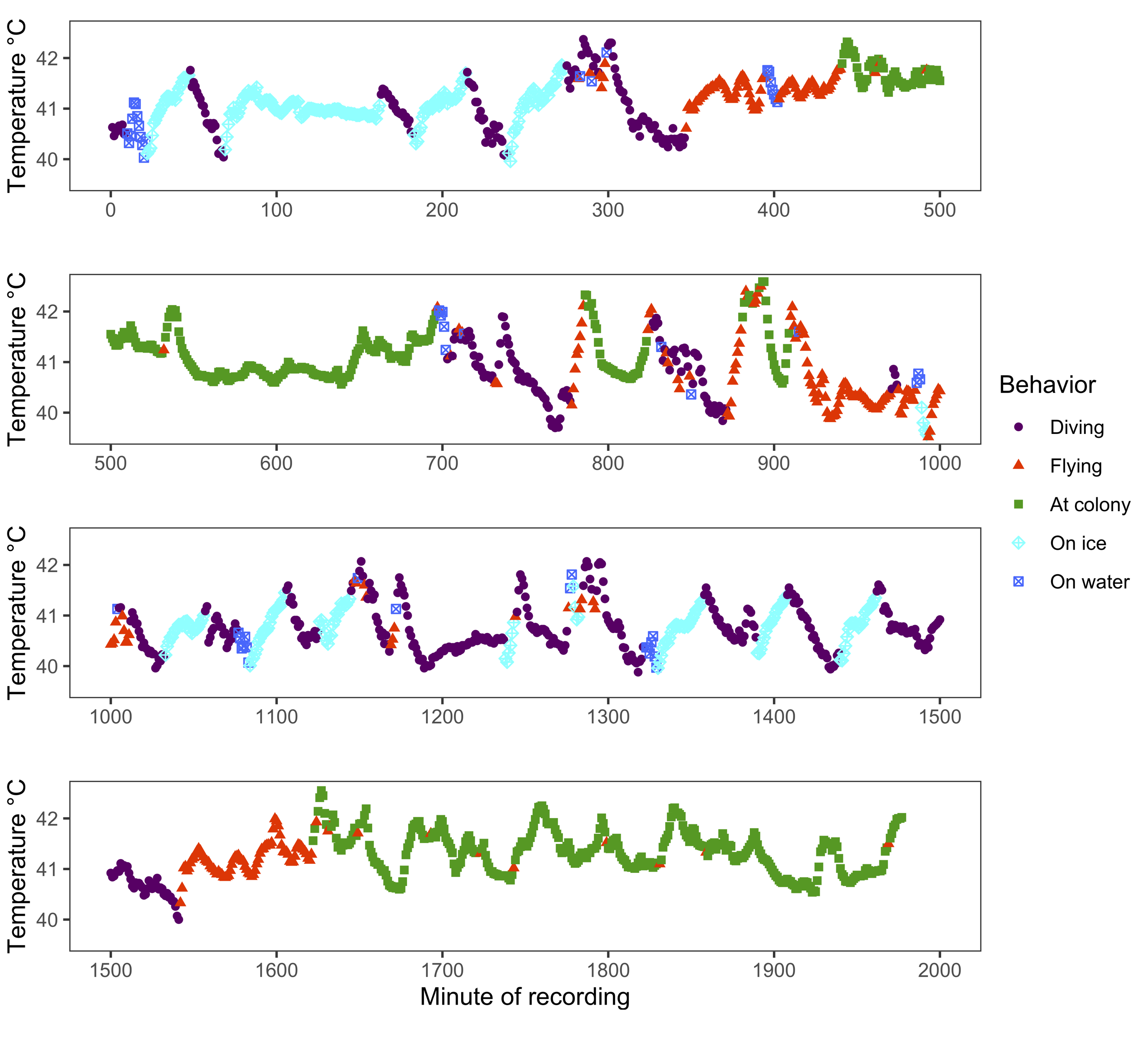

Fig. S3. Variation in body temperature (T_b_**)** through time and color coded with respect to behavioral state for one individual little auk (LIAK20EG14) from the Ukaleqarteq, East Greenland, population. Note the rebound in T_b_ on sea ice following declines during diving. T_b_ can be observed to increase while flying in several cases, but again T_b_ appears to decrease in flight after leaving the colony, which may reflect departure from a thermal refuge (see also Fig. S1). In this individual, we observe some variability in T_b_ at the colony, associated with interspersion of short flying episodes. Little auks often take flight at the colony when threatened by predators (especially glaucous gulls (*Larus hyperboreus*)), and associated activation of stress responses may result in elevation of, and variation in, T_b._

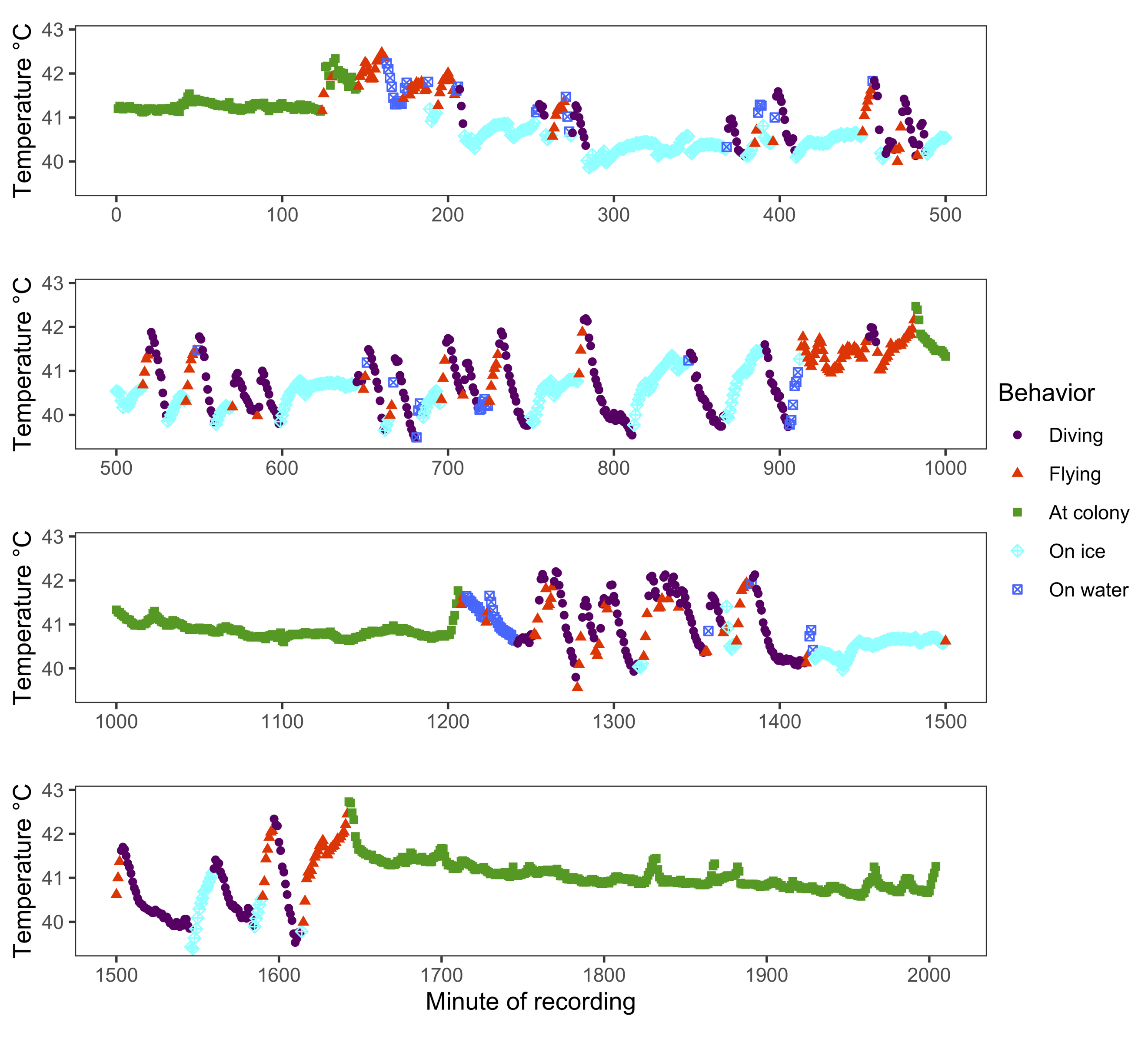

Fig. S4. Variation in body temperature (T_b_**)** through time and color coded with respect to behavioral state for one individual little auk (LIAK20EG17) from the Ukaleqarteq, East Greenland, population. Note rebounds in T_b_ when on the sea ice following declines while diving in cold polar waters. T_b_  can also be observed to increase with time during flight and decline with time at the colony. Overall stability of T_b_ at the colony is also apparent.

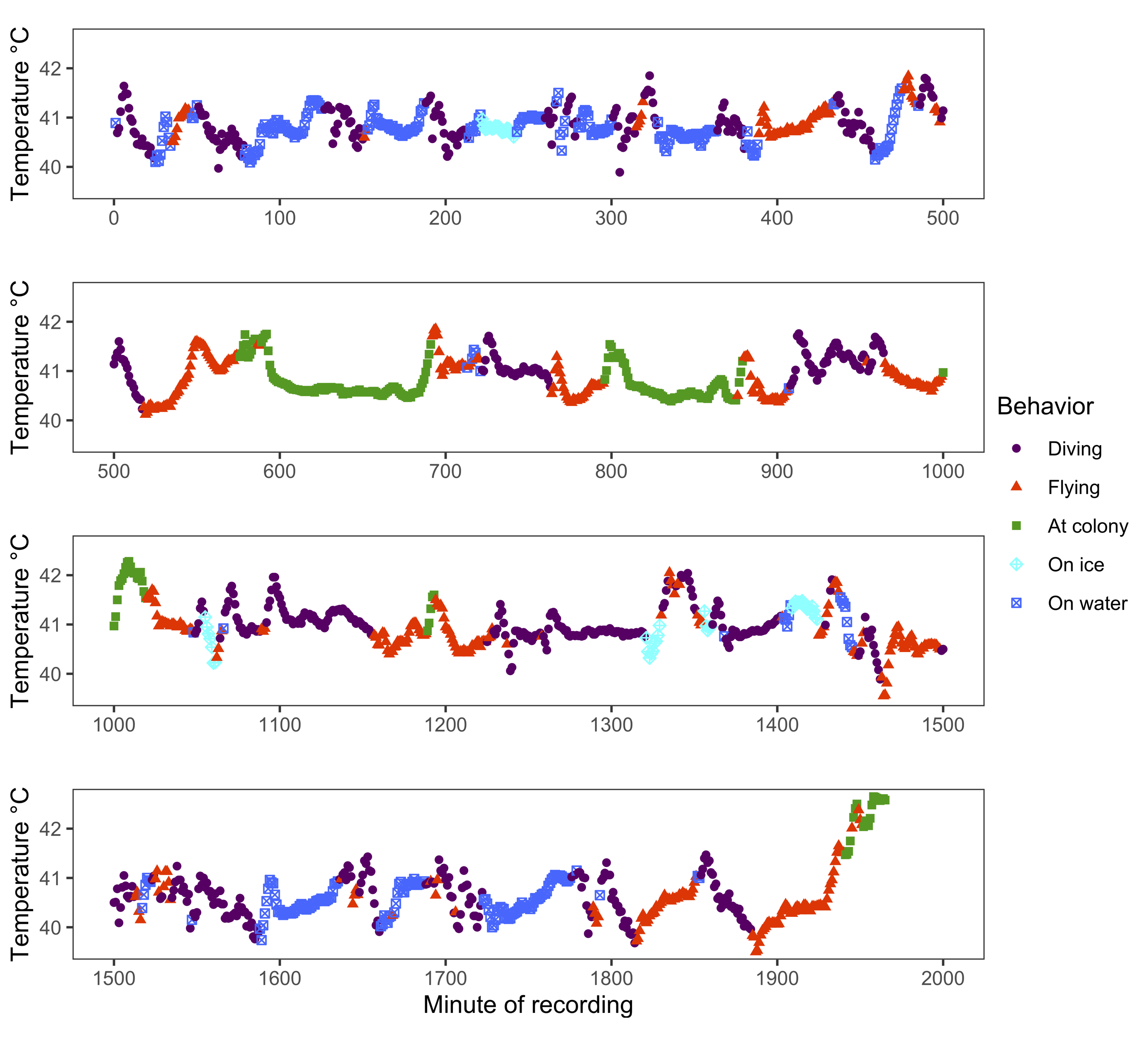

Fig. S5. Variation in body temperature (T_b_**)** through time and color coded with respect to behavioral state for one individual little auk (LIAK20EG24) from the Ukaleqarteq, East Greenland, population. This bird spent little time on ice, and the rebound in T_b_ on ice following diving bouts is not clear. T_b_ can be observed to increase during flight in several instances, and to decline at the colony. While resting on the water surface, the pattern of change in T_b_ with time appears variable.

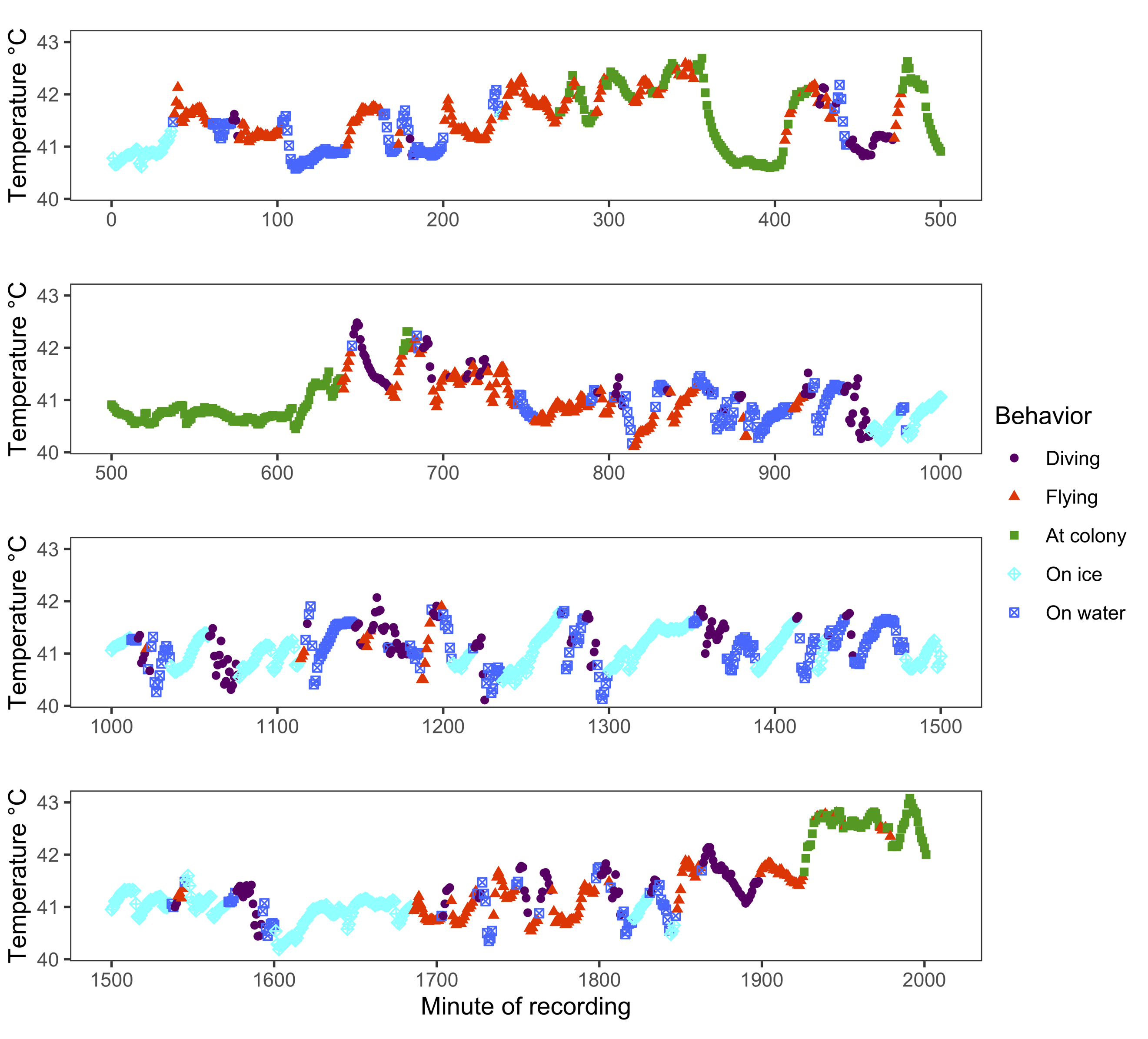

Fig. S7. Variation in body temperature (T_b_**)** through time and color coded with respect to behavioral state for one individual little auk (LIAK20EG25) from the Ukaleqarteq, East Greenland, population. In this bird, the rebound in T_b_ on ice following foraging bouts (diving) is apparent. T_b_ can also be observed to increase during flight in serval instances. Interspersion of short flight episodes with time at the colony can be observed to increase variability of T_b_ at the colony (e.g. ~minute 275-350).

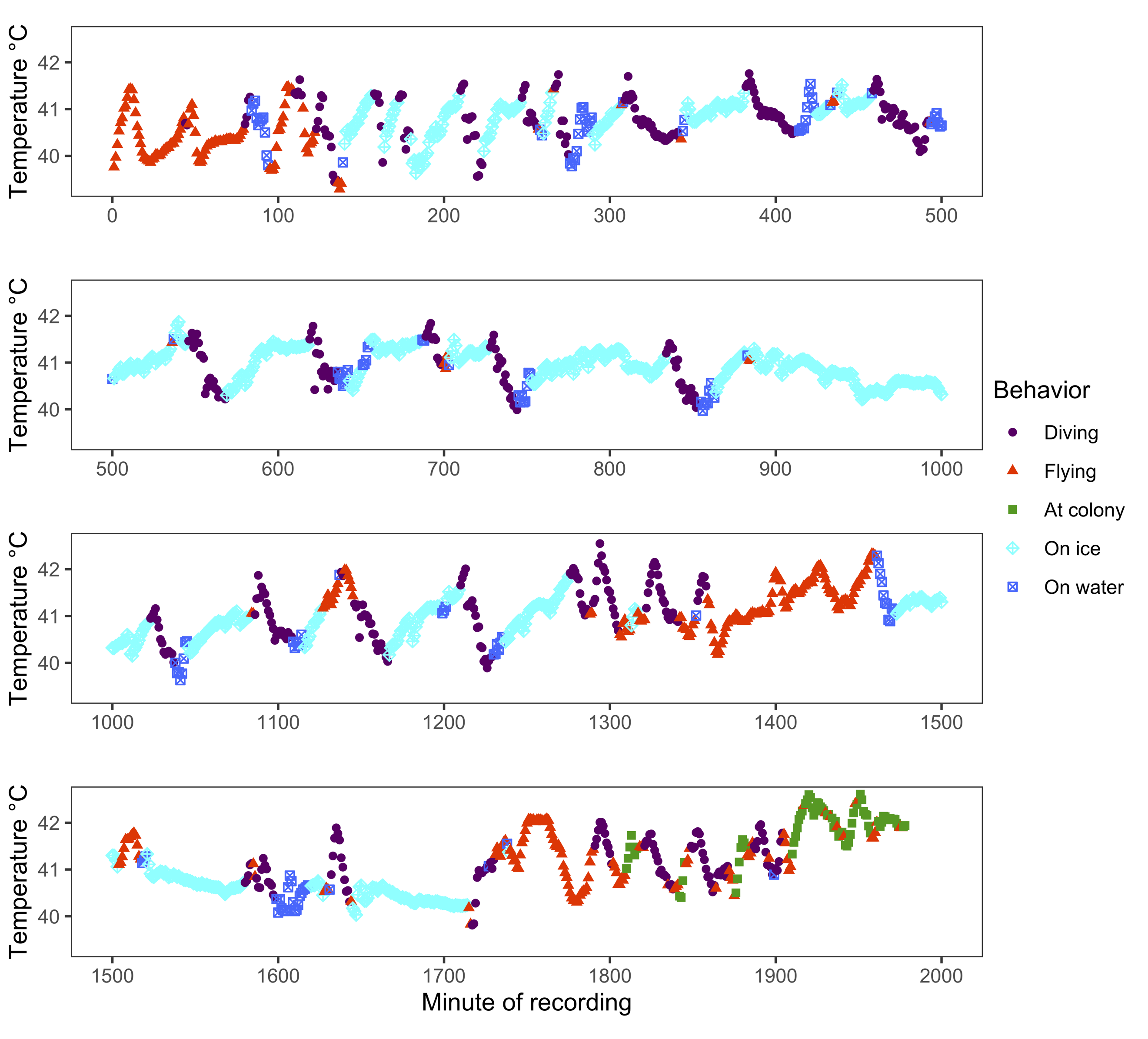

Fig. S8. Variation in body temperature (T_b_**)** through time and color coded with respect to behavioral state for one individual little auk (LIAK20EG28) from the Ukaleqarteq, East Greenland, population. The rebound of T_b_ on ice following foraging bouts (diving) is apparent. In this individual the pattern of T_b_ variation through time in flight shows some increases and decreases. The decreases are somewhat difficult to explain in these cases, but could reflect changes in exposure to wind.
